# Modelling the interplay between the CD4^+^/CD8^+^ T-cell ratio and the expression of MHC-I in tumours

**DOI:** 10.1101/2020.07.03.185926

**Authors:** Christian John Hurry, Alexander Mozeika, Alessia Annibale

**Affiliations:** Department of Mathematics, King’s College London, Strand, London WC2R 2LS, United Kingdom; London Institute for Mathematical Sciences, 35A South Street, London, W1K 2XF, United Kingdom; Institute for Mathematical & Molecular Biomedicine, King’s College London, Hodgkin Building, London, SE1 1UL, United Kingdom

**Keywords:** CD4/CD8 ratio, MHC-I, immunology, statistical mechanics

## Abstract

Describing the anti-tumour immune response as a series of cellular kinetic reactions from known immunological mechanisms, we create a mathematical model that shows the CD4^+^/CD8^+^ T-cell ratio, T-cell infiltration and the expression of MHC-I to be interacting factors in tumour elimination. Methods from dynamical systems theory and non-equilibrium statistical mechanics are used to model the T-cell dependent anti-tumour immune response. Our model predicts a critical level of MHC-I expression which determines whether or not the tumour escapes the immune response. This critical level of MHC-I depends on the helper/cytotoxic T-cell ratio. However, our model also suggests that the immune system is robust against small changes in this ratio. We also find that T-cell infiltration and the specificity of the intra-tumour TCR repertoire will affect the critical MHC-I expression. Our work suggests that the functional form of the time evolution of MHC-I expression may explain the qualitative behaviour of tumour growth seen in patients.

**Mathematics Subject Classification (2010):** MSC 37C25 · MSC 82C99 · MSC 37N25 · MSC 92B99

## 1 Introduction

It is now known that tumours evoke an immune response, which can alter the growth and makeup of a tumour. Greater understanding of the anti-tumour immune response has led to the development of immunotherapies, which have seen some degree of success in clinical trials, particularly in cancers of the blood (Kochenderfer et al, 2014; Brentjens et al, 2013; Leach et al, 1996; McDermott et al, 2015; Hodi et al, 2010). However, the use of these therapies in solid cancers and conversion to the clinic still remains a challenge as was pointed out by Kakarla and Gottschalk (2014).

The immune system can be seen as a network of interacting cells, with many different cell types working together to perform a wide-ranging and robust function against pathogens. In addition to this, tumours are rapidly evolving parts of this network. A systemic understanding of the mechanisms of the anti-tumour immune response is, therefore, vital in the development of immunotherapies. Immediately, this questions what should be considered important systemic processes and what can be regarded as non-dominant behaviour. With this in mind this paper seeks to address the adaptive, T-cell dependent anti-tumour immune response.

T-cells are lymphocytes, a group of white blood cells, distinguished from other lymphocytes by their unique receptor known as the T-cell receptor (TCR). A TCR is specific to a particular antigen (a part of a protein recognised by a TCR). In addition to this, T-cells can be split into two main sub-types: helper and cytotoxic T-cells. Antigens are picked up by professional antigen presenting cells (APCs) and presented as a peptide on the surface molecule MHC-II. Helper T-cells can then activate by binding to MHC-II which displays the antigen that the TCR is specific to. Activated helpers can then activate cytotoxic T-cells which eliminate cells that present their conjugate antigen via MHC-I. The T-cell dependent response eliminates infected cells, but is also invoked by tumours (Restifo et al, 2012).

A common experimental technique in immunology is immunostaining, which uses antibodies that bind to specific proteins, to sort cells. Helper T-cells are known for their high expression of the protein CD4, and cytotoxic cells for high CD8. For this reason, they are commonly referred to as CD4^+^ and CD8^+^ cells, respectively. A decrease in the CD4^+^/CD8^+^ ratio is considered to be a good prognostic marker for conditions associated with immunodeficiency such as HIV (Taylor et al, 1989; Serrano-Villar et al, 2014), and aging (Wikby et al, 1998; Olsson et al, 2001).

Solid tumours are a porous mixture of tumour cells, immune cells and healthy tissue cells. T-cells can infiltrate tumour cells and the CD4^+^/CD8^+^ ratio of infiltrating T-cells can be measured from tumour samples. A low CD4^+^/CD8^+^ of infiltrating T-cells has been considered as a marker for prognosis across several cancer types including: cervical (Sheu et al, 1999), breast (Sheu et al, 2008; Sevcíková et al, 1992), lung, liver, testicular and colorectal cancers (Tancinia et al, 1990). Tancinia et al (1990) surveyed tumours from a cohort of breast, lung, colorectal, liver and testicular cancers, and found that low intra-epithelial CD4^+^/CD8^+^ was associated with early stage cancer due to an expanded CD8^+^ population, and that later stage cancers were associated with low CD4^+^/CD8^+^ due to a loss of the CD4^+^ population. In addition to these results in solid cancers, a low CD4^+^/CD8^+^ ratio in the blood correlated with poor survival in chronic lymphocytic leukemia (CLL) (Nunes et al, 2012). Despite the evidence that across cancer types a low CD4^+^/CD8^+^ ratio is a sign of poor prognosis, patients with low CD4^+^/CD8^+^ ratio of tumour infiltrating T-cells were found to have significantly improved survival in separate studies of colorectal cancer (Diederichsen et al, 2003) and ovarian cancer (Sato et al, 2005), contradicting these studies.

There are many factors, biological and immunological, which could account for the prognostic variability in the CD4^+^/CD8^+^ ratio. Overall T-cell infiltration, i.e the density of T-cells in a tumour, may account for this variability in tumours, as the absolute number of T-cells will affect the strength of the immune response. As one would expect, the density of T-cells has also been shown to correlate with prognosis in cancer (Ménard et al, 1997; Ryschich et al, 2005). However, recent results have shown that the tumour reactivity of infiltrating T-cells, which reflects the proportion of TCRs that are specific to tumour associated antigens, is low in cancers where infiltration is a marker for prognosis (Scheper et al, 2019). Therefore, the infiltration of specific T-cells may also lead to variability in prognostic markers.

A key stage in the progression of tumours is the down-expression of MHC-I, a cell surface molecule with which immune cells interact (Garcia-Lora et al, 2003). The effect of this on tumour growth has been reportedly mixed. The total absence of MHC-I in breast tumours has been shown to activate a group of innate immune cells, natural killer (NK) cells, which can kill cells without MHC-I recognition (Madjd et al, 2005). However, studies of colorectal cancer showed that tumours with high MHC-I were found in patients with longer survival times and that MHC-I could be used as an independent marker of prognosis (Watson et al, 2006; Simpson et al, 2010). To complement the results seen with breast cancer, the total absence of MHC-I in colorectal cancer also showed longer patient survival times compared with low MHC-I expression, due to the activation of NK cells (Watson et al, 2006). To summarise, MHC-I expression impacts the growth of tumours *in vivo*, with low expression favouring tumour growth, while high expression leading to longer patient survival. The exception is tumours with an absence of MHC-I which trigger an innate immune response from NK cells. This stresses the importance of tumour heterogeneity: low MHC-I expression, due to a large proportion of cells down-expressing MHC-I, will prevent a sufficient T-cell response, but the low level of MHC-I will also interrupt the response from NK cells.

There is a mechanistic interplay between the CD4^+^/CD8^+^ ratio and MHC-I expression. This is because cytotoxic cells, which form the majority of CD8^+^ cells, bind to MHC-I in order to eliminate tumour cells. In spite of this fact there is a lack of data measuring the CD4^+^/CD8^+^ ratio and MHC-I together. One study has shown that the prognostic value of T-cell markers improves when MHC-I expression is also considered (Turcotte et al, 2014). Additionally, the loss of MHC-I in pancreatic cancer has been shown to lead to a lower level of infiltration by cytotoxic T-cells (Ryschich et al, 2005). Cytotoxic T-cell infiltration was found to be a marker of prognosis, however, MHC-I alone was not. Our hypothesis is that the interplay between CD4^+^/CD8^+^, T-cell infiltration, and MHC-I could explain the differences in prognostic value for these parameters across individual tumours and different cancers. To address this we create a mathematical model, starting from known cellular processes to derive system-wide behaviour. The aim is to see if such a model captures this interplay and explains potential variation in the prognostic value of CD4^+^/CD8^+^ and MHC-I.

As research has turned towards systemic modelling of the immune system there has been an increase in mathematical and computational approaches to modelling challenges in immunology. Mathematical models used are mainly *deterministic*, comprised of ordinary differential equations (ODEs). Such models usually fall into two extremes: either, the model is low dimensional, ‘macroscopic’, and does not fully capture the systemic behaviour of the system; or it is high dimensional, ‘microscopic’, with a large number of unknown parameters; usually this makes statistical evaluation of these models with real data difficult. Furthermore, ODE models also fail to capture the inherent *stochasticity* of biological processes. In this paper we consider a model that lies between these two approaches, keeping the model to a few key parameters, whilst also including inherent stochasticity of microscopic behaviour.

The cytotoxic anti-tumour response has been modelled previously, most notably by Kuznetsov et al (1994). Despite their pioneering efforts, they neglected to include the immunosupressive effects of the tumour microenvironment, something which was later accommodated for by Dritschel et al (2018) who included specifically the interactions between the helper and cytotoxic T-cell populations. Both models are deterministic and our work deviates from them in this respect.

In our work we model the interplay between the CD4^+^/CD8^+^ T-cell ratio and the expression of MHC-I in tumours explicitly. Although our model is also a system of ODEs which describe the change in concentration of cells, our model differs from previous efforts by including the stochastic dynamics of T-cell activation. The latter is used in the derivation of our model, but our analysis is based on non-equilibrium statistical mechanics, as similarly implemented by Annibale et al (2018), allowing us to describe the macroscopic behaviour of the system deterministically. Statistical mechanics has a rich recent history in modelling the immune system (Perelson and Weisbuch, 1997; Lucia and Maino, 2002; Chakraborty and Košmrlj, 2010; Mora et al, 2010; Agliari et al, 2013; Bartolucci et al, 2016) and here we use it to address the unique modelling challenges that tumour immunology poses. In doing so we derive a condition for the eradication of tumours which reveals a critical threshold of MHC-I expression. This threshold is found to depend on the CD4^+^/CD8^+^ T-cell ratio, T-cell infiltration, and T-cell specificity.

In this work, we exclude the effects of NK cells and macrophages, focusing exclusively on the T-cell dependent response. This allows our model to be kept to a few key parameters that can be analysed in full. As previously discussed, NK cells play a dominant role when MHC-I is totally *absent*, a case which is less relevant for our model. The role of macrophages in the tumour is a double edged sword, with macrophages eliminating tumour cells, and also forming part of the bulk tumour. This behaviour is complex, and has been modelled elsewhere (Eftimie and Eftimie, 2018), and is unlikely to affect our analysis on the interplay between the CD4^+^/CD8^+^ T-cell ratio and MHC-I. The result of this work is a low dimensional model with just a few important parameters, derived from known immunological and biological mechanisms.

The remaining sections of this paper will be organised as follows: in Sec. 2 we will introduce our mathematical model for tumour-immune interactions, in Sec. 3 we will present four key results that are of biological relevance and we will provide a discussion that assesses how our results fit in with the questions raised by the literature. We will discuss benefits and limitations of our modelling approach as well as pathways for future work in Sec. 4. Technical details are described in the appendix.

## 2 Constructing a mathematical model

### 2.1 The adaptive anti-tumour immune response

The anti-tumour immune response can be described by a series of cellular kinetic rate reactions. Here we list the full set of reactions that we model. By describing the immune response in this way, we can use the law of mass action to write a series of ODEs for further analysis. We consider the anti-tumour immune response to be between tumour cells, antigen presenting cells and T-cells, which are split into two sub-types, helper and cytotoxic. Their interactions, summarised in figure 1, are governed by the following reactions:

**Fig. 1.**
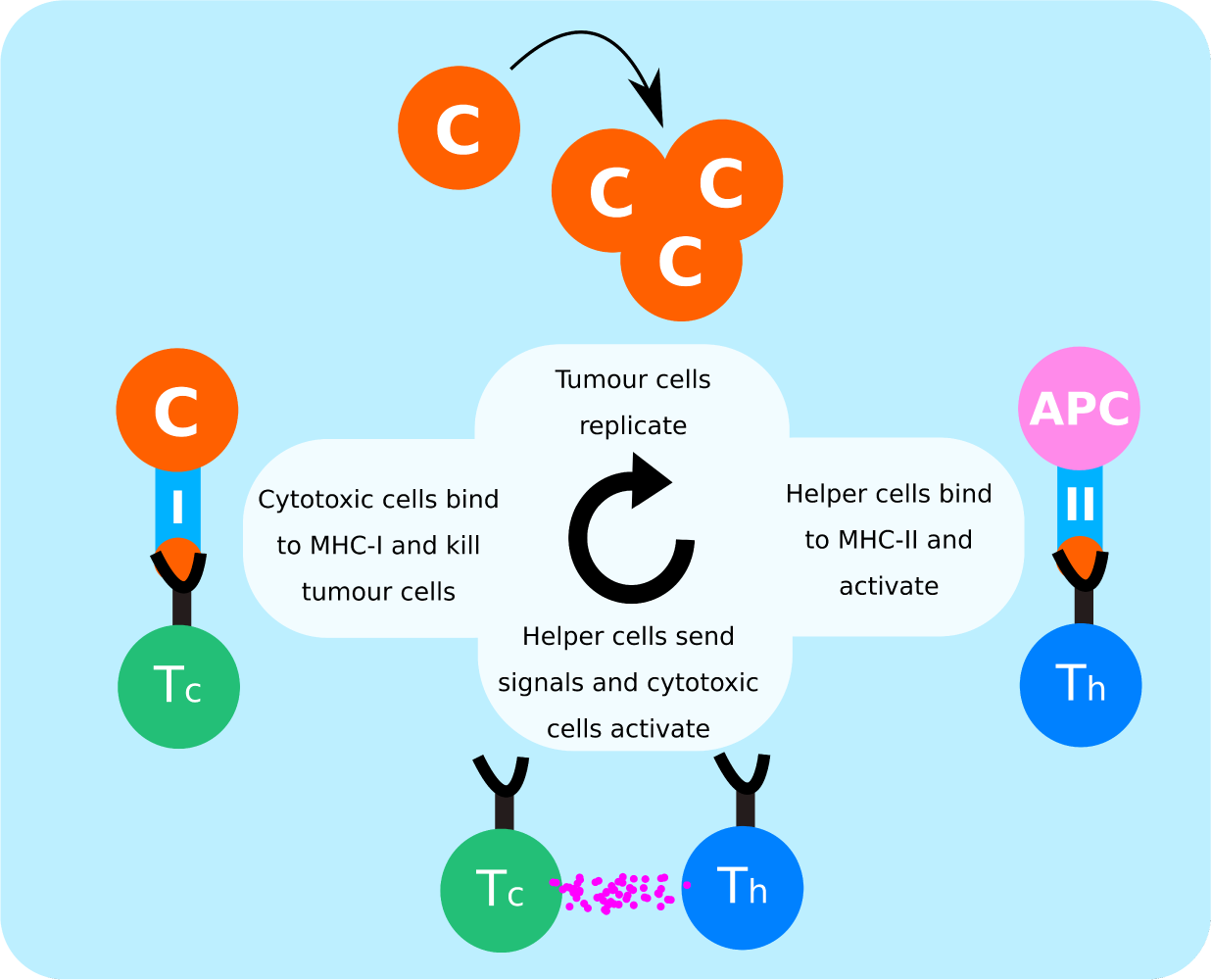
Summary of cellular interactions between tumour cells (C), antigen presenting cells (APCs), helper T cells (Th) and cytotoxic T cells (Tc) in anti-tumour immune response

- Tumour cells, *C*, replicate at a constant rate *r* and produce free floating antigens that are detected by the immune system,

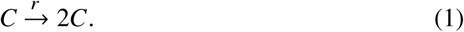
- Tumour cells compete for resources such that there is a maximum concentration of tumour cells, *ρ*_*c*_, in a finite volume

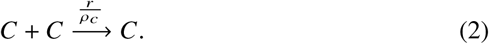
- Professional antigen presenting cells, *APC*, such as dendritic cells, take up free floating antigens, process them into peptides, and display them on surface molecules MHC-II, *APC*\*,

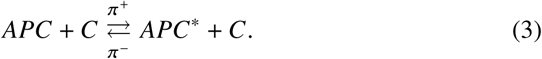
- Helper T-cells, *T*_*h*_, bind to the peptide-MHC-II complex and activate, 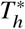,

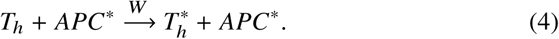
- Cytotoxic T-cells, *T*_*c*_, are activated, 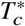, by both cytokines from activated helpers and stimulation from tumour cells that the cytotoxic T-cells are specific to,

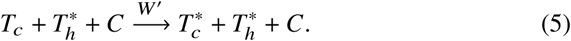
- Activated helpers induce the proliferation of antigen presenting cells

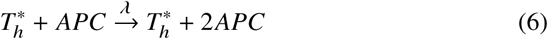
- Activated cytotoxic T-cells eliminate tumour cells after docking to MHC-I,

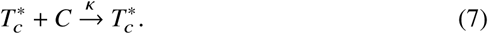

The ‘killing’ rate, κ, is proportional to the rate at which cytotoxic cells bind and induce cytotoxic death, *k*, and the expression of MHC-I, *γ*, which can decrease throughout the development of the tumour. This is such that κ = *kγ*. For the remainder of this paper we set *k* = 1 and, without loss of generality, focus on the expression of MHC-I, *γ* ∈ [0, 1], with *γ* = 0 corresponding to no tumour cells expressing MHC-I and *γ* = 1 corresponding to maximum MHC-I expression in the tumour. In principle, *γ* will vary with time, but to simplify the analysis we assume for now that it is constant and comment later on the effect of time-dependent *γ*.

### 2.2 A statistical mechanics description of the immune system

We consider a solid tumour to occupy a space of fixed volume *V*. This volume is comprised of cells including helper and cytotoxic T-cells, professional antigen presenting cells and tumour cells. The volume is permeable such that T-cells can move freely from the periphery into the solid tumour. We consider there to be a large number, *N*, of T-cells such that the density of T-cells, 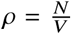, is finite. T-cells are a heterogeneous population of cells, so to model this we describe each T-cell by three random variables *σ*_*i*_, *η*_*i*_, *ξ*_*i*_ which respectively describe the activation, sub-type and specificity of a T-cell. Describing T-cell characteristics with random variables is a modelling choice that will make use of methods of statistical physics. By doing so, we can evaluate our results for different populations of T-cells with different proportions of sub-type and specificity. The sub-type of the T-cell is described by the random variable *η*_*i*_,

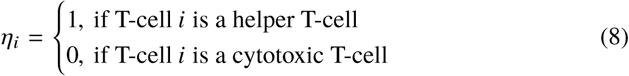

drawn from the distribution,

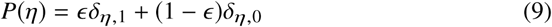

such that the parameter *ϵ* controls the proportion of helper and cytotoxic T-cells. The parameter *ϵ* takes values in the range [0, 1] such that if *ϵ* = 0 all T-cells are cytotoxic and if *ϵ* = 1 all T-cells are helpers. The helper/cytotoxic ratio is given by 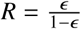.

Each T-cell has receptors known as T-cell receptors (TCRs). A given TCR is specific to an antigen, and if they are not specific to tumour associated antigens, they will not form part of the immune response. To model this we define the random variable *ξ*_*i*_ such that

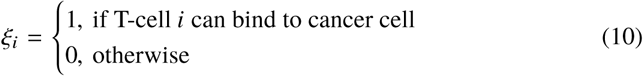

which is drawn from the distribution,

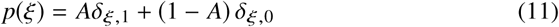

such that *A* ∈ [0, 1] controls the fraction of T-cells that are specific to tumour associated antigens. To simulate a heterogeneous population of T-cells we draw *N* random numbers from each of the above distributions. Finally, activation of each T-cell is described by the time dependent state variable *σ*_*i*_(*t*) where,

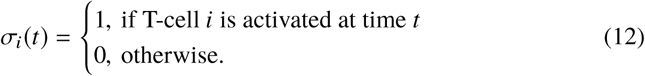

The activation of each T-cell evolves with time over the course of the immune response and we will later define the stochastic dynamics of T-cell activation.

With this description of the microscopic detail of each T-cell in the system, we can then proceed to write ordinary differential equations, modelling the cellular reactions in equations (1)-(7), which describe the concentration of the other cells in the system. The number of tumour cells, antigen presenting cells, and antigen presenting cells with MHC-II peptide complex are represented by the variables [*C*], [*B*] and [*P*], respectively. Their concentration of each of these cells is then simply given by their number over the volume of the solid tumour, *c* = [*C*] /*V, b* = [*B*]/ *V* and *p* = [*P*]/ *V*. The evolution of the concentration of each of these cells can be written in the following way

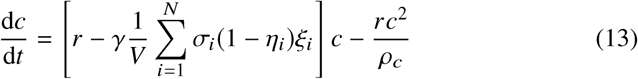

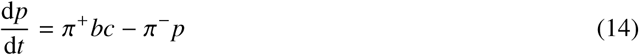

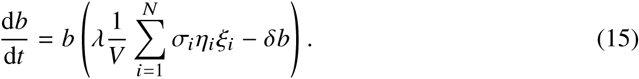

Equation (13) contains three terms describing the change in tumour cell concentration with time, 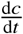. The first term states that *c* will increase at a rate *r* proportionally with *c*, whereas the third term states that tumour cells will compete for resources and reach a carrying capacity concentration *ρ*_*c*_. The second term describes the effect of the T-cells on the tumour cells: a sum is taking over all the T-cells, and a non-zero contribution is only made by T-cells which are cytotoxic, specific to tumour cells and active, i.e. when *σ*_*i*_ = 1 − *η*_*i*_ = *ξ*_*i*_ = 1. Therefore, the second term states that tumour cells will be killed at a rate *κ* = *γ* and proportionally with the concentration of tumour cells and the fraction of active, specific cytotoxic T-cells. The equations for 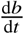 and 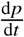 can similarly be annotated, and will match the cellular reactions previously described. In principle, equation (15) should contain a term reflecting the loss of naive APCs becoming activated APCs, 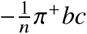, and a gain term reflecting the reverse process 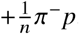. Here the *n* represents the load of APCs, the number of antigens that they can process at one time. Since APCs can process multiple antigens via multiple pathways, we consider *n* to be large such that these terms are negligible.

From a set of cellular reactions we have started to build a set of ordinary differential equations, with the inclusion of a microscopic description of T-cells. As they stand we can not solve equations (13)-(15) directly without prescribing dynamics for the activation variables *σ*_*i*_. In order to implement the latter we define the macroscopic *observables*,

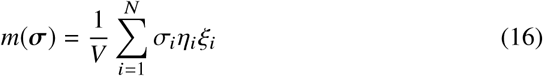

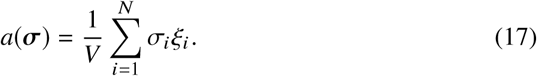

where ***σ*** ∈ {0, 1} ^*N*^, representing the density of active, specific helpers and the density of active, specific T-cells, respectively. With these definitions equations (13)-(15) can be written as follows,

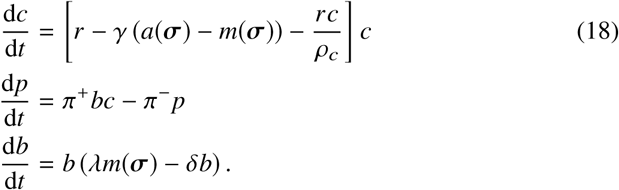

We now seek to derive the time evolution of the macroscopic observables *a*(***σ***) and *m*(***σ***) in order to get a closed system of ODEs.

### 2.3 Macroscopic dynamics of T-cell activation

To close the system of equations (18), we must define the dynamics of T-cell activation ***σ***(*t*). During the immune response, T-cell activation occurs through a TCR-dependent pathway whereby the TCR binds to the peptide complex of MHC-II and a condition is met such that the T-cell activates. The precise nature of the activation of T-cells is debated, but there is evidence to suggest that sufficient binding time is required for activation (Allard et al, 2017; Tian et al, 2007; Robert et al, 2012; Tkach and Altan-Bonnet, 2013; Aleksic et al, 2010). Due to noise, inherent in biological systems, T-cells are likely to bind and unbind in a stochastic manner, therefore we treat T-cell activation as a stochastic process. For simplicity, we assume that T cells update their activation state at regular time intervals of duration Δ (which will eventually be sent to zero to retrieve the continuous time dynamics), according to the rule

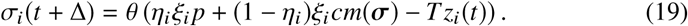

Here *z*_*i*_(*t*) is a random “activation” noise and the “temperature” parameter *T* controls its magnitude. Also, we have used the Heaviside step function, θ (*x*), such that θ*x* = 1 for *x* > 0 and 0 otherwise. For a T-cell to be active at time *t* + Δ, the argument of θ must be positive. If we consider no noise, i.e *T* = 0, and consider helper T-cell *i*, i.e *η*_*i*_ = 1, only the first term, *η*_*i*_*ξ*_*i*_ *p*, inside θ remains. Therefore, helper T-cell *i* will activate if it is specific, i.e *ξ*_*i*_ = 1, and there are antigen presenting cells with tumour associated antigens, *p* > 0. If T-cell *i* is cytotoxic, i.e *η*_*i*_ = 0, then the only non-zero term is (1 − *η*_*i*_) *ξ*_*i*_ *cm*(***σ***) such that activation requires the presence of tumour cells, *c* > 0, and active helpers, *m*(***σ***) > 0. Noise is introduced when *T* > 0 and can be interpreted as the amount that T-cells deviate from their deterministic activation rules, thus modelling stochasticity of T-cell activation. In addition to the binding and unbinding of T-cells to MHC-I, noise also accounts implicitly for alternative activation pathways. For example, cytotoxic T-cells may not necessarily need helper T-cells to activate, they can be directly activated by APCs. Additionally, immunosuppressive effects (e.g regulatory T-cells) not explicitly considered here are implicitly accounted for in this noise term.

Next, one must specify the statistical properties of the activation noise. Information about the latter is very scarce in biological literature (Irvine et al, 2002; Wedagedera and Burroughs, 2006; Burroughs and Van Der Merwe, 2007) and a natural choice would be to assume a Gaussian distribution of noise but, for convenience of analysis and without loss of generality, we consider a distribution of noise more common in statistical physics (see Appendix for details).

Assuming that T cells are updated sequentially, equation (19) can be cast into a master equation for the time-dependent probability *P*_*t*_ (***σ***), by setting Δ = 1 / *N* in equation (19), and taking the limit *N*→ ∞. This is done in the Appendix where equations for the time evolution of the macroscopic observables *a*(***σ***) and *m*(***σ***) are also derived. It will turn out that fluctuations of these quantities around their averages, *m*(*t*) = Σ ***σ*** *P*_*t*_ (***σ***) *m*(***σ***) and *a*(*t*) = Σ ***σ*** *P*_*t*_ (***σ***) *a*(***σ***), vanish as the number of T-cells, *N*, is sent to infinity and that the evolution of these averages is governed by the equations

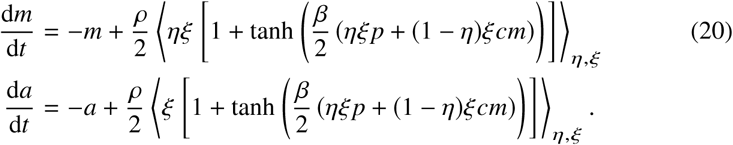

In the above the density of T-cells 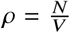 is assumed to be finite when *N* → ∞ and ⟨… ⟩ _*η,ξ*_ denotes the average over the joint distribution,

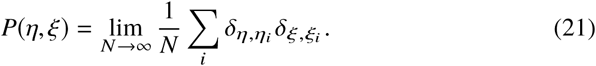

Finally, assuming that *P*(*η*, *ξ*) = *P*(*η*) *P*(*ξ*), i.e. the ability of a T cell *i* to bind to cancer does not depend on whether *i* is helper or cytotoxic, and using (9) and (11) to compute the averages in (20), we obtain the closed system of equations,

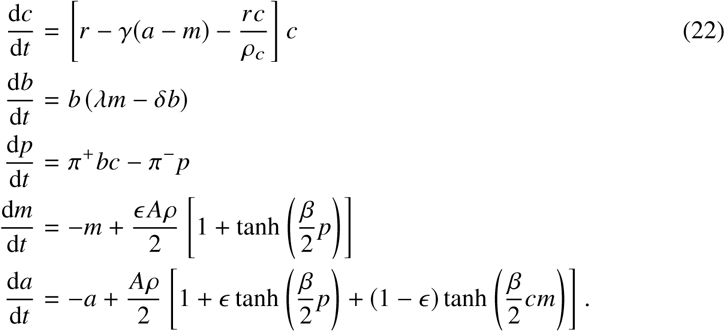

## 3 Results

### 3.1 Conditions for tumour eradication

We use the system of equations (22) to derive a set of conditions which will qualitatively describe how the anti-tumour immune response changes with parameters of the model. First, we write the system of equations in a more compact way by defining the vector **x** = (*c, b, p, m, a*) such that 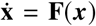, where each component of the vector **F** is the RHS of the corresponding ODE. Second, we find fixed points of the dynamics from 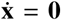 and analyse their stability by inspecting the eigenvalues of the Jacobian ∂/**F**∂**x**.

We find that there are two fixed points which subject to some condition are stable. There are two other fixed points but they are always unstable. The potentially stable fixed points are given by,

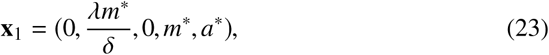

where 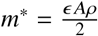 and 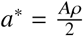 and

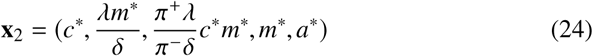

where *c*\*, *m*\* and *a*\* are the solution to the system of equations

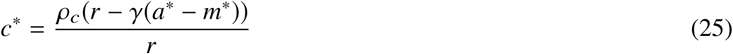

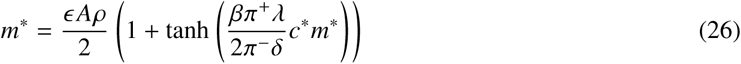

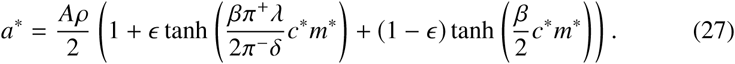

**x**_1_ corresponds to tumour eradication, *c*\* = 0, whereas **x**_2_ corresponds to tumour escape, *c*\* ≠ 0. The size of the tumour at **x**_2_ varies with parameters of the system. When T-cells are all cytotoxic, *ϵ* = 0, there is no signal from helper cells *m*\* = 0, and all T-cell signal comes from cytotoxic cells, 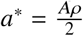, resulting in a tumour below the carrying capacity, 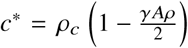. However, when all T-cells are helpers, *ϵ* = 1, we have that the net T-cell signal is equivalent to the helper T-cell signal, *m*\* = *a*\*, and that the tumour reaches the carrying capacity, *c*\* = *ρ*_*c*_.

From analysis of the eigenvalues of the Jacobian ∂/**F**∂**x** we find that **x**_1_ is stable when,

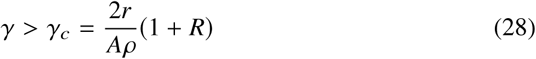

when this condition is not met **x**_1_ is unstable. To assess the stability of **x**_2_ we should analyse the eigenvalues of the Jacobian evaluated at **x**_2_, however the eigenvalues are found to be non-trivial and a condition for stability based on a single parameter as in (28) is not tractable.

To make progress analytically, we look at the long-time dynamics of the model. To this end, we separate three different timescales of immune response dynamics: the timescale of tumour cell division *τ*_*c*_, antigen presentation *τ*_*p*_, and T-cell activation *τ*_*a*_. This implies that time should be a re-scaled in (22) as follows,

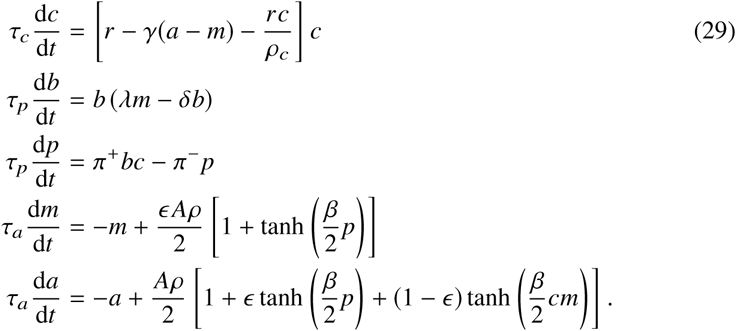

To study the long time dynamics the relative magnitude of these timescales must be specified. We assume that *τ*_*p*_ → 0 such that the processing and presentation of antigens is fast implying that the variables changing at this timescale will approach their stable nullcline, 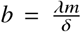 and 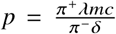. Then there are two options: either tumour cell division is a faster process than T-cell activation or vice versa. If we consider the case that T-cell activation is faster than tumour cell division, i.e *τ*_*a*_ ≪ *τ*_*c*_, then we can formally send *τ*_*a*_ → 0 and find that the T-cell activation variables approach their nullclines,

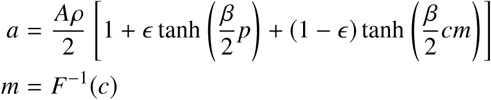

where

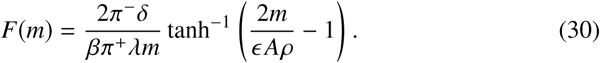

This reduces the system (29) to a single ODE,

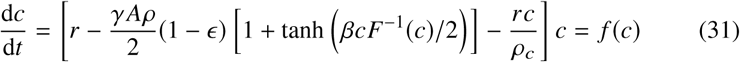

which has two fixed points given by

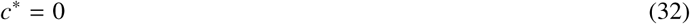

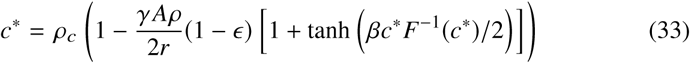

corresponding, respectively, to tumour eradication and large stable tumour formation. The non-trivial fixed point, *c*\* > 0, can be found using the relation *F*^−1^ (*c*\*) = *m*\* via the numerical solution of (25)-(27).

To inspect the stability of the fixed points we evaluate the derivative *f* ′ (*c*) of the ‘velocity’ function defined in (31). The fixed point *c*\* = 0 is stable when *f*′(0) < 0 giving us the condition 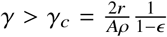 which is equivalent to (28). The non-trivial fixed point *c*\* < 0 will be stable when *f*′ (*c*\*) <0, yielding

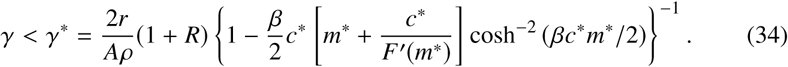

Since it is not necessary that *γ*_*c*_ = *γ*\* there is a possibility that either i) *γ*\* < *γ* < *γ*_*c*_ and both fixed points are unstable or ii) *γ*_*c*_ < *γ* < *γ*\* and both fixed points are stable; this would suggest that the dynamics are non-trivial and can not be analysed through linear stability analysis alone. To investigate which, if any, of the two scenarios is taking place, we perform limiting analysis of the stability condition. We first note that for meaningful values of *m* i.e 0 < *m* < *ϵ* A*ρ* which must be, by definition, at most of order *𝒪* (1), we can show that *F*′(*m*) > 0. The term 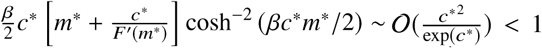 which vanishes at both *c*\* → 0 and *c*\* → ∞. It then follows from equations (33) and (34) that,

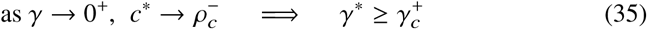

with equality at *ρ*_*c*_ → ∞. We can also look at the limit that the *γ* approaches its critical value to find,

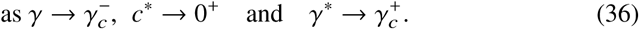

However, the exact value of *γ*\* can only be found through numerical solution of equations (25) and (26). Solving the latter, see figure 2 (left panel), we find that

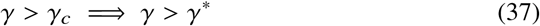

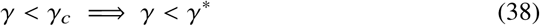

which implies that the fixed points exchange stability as *γ* →*γ*_*c*_.

**Fig. 2.**
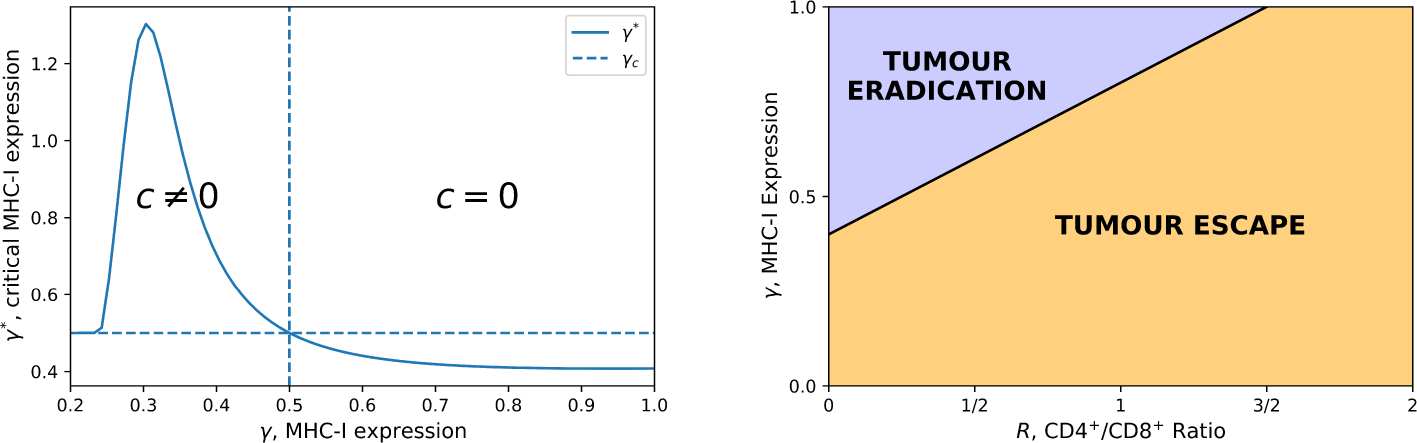
Left: The fixed point of equation (31) where *c* ≠ 0 is stable when *γ* < *γ*\*. We solve equation (33) numerically to plot *γ*\* as a function of *γ*. When *γ* > *γ* _*c*_ the fixed point where *c* = 0 is stable. The value of *γ*_*c*_ is indicated by the dashed lines. The model parameters used are: tumour replication rate *r* = 0.15, CD4/CD8 ratio *R* = 1 3, and specific T-cell density *Aρ* = 0.8. All other parameters are set to 1. Right: *γ*_*c*_ is plotted against the helper/cytotoxic ratio *R*. Tumour cells are removed above the critical line. The model parameters used are: tumour replication rate *r* = 0.2, specificity of T-cells *A* = 1 and density of T-cells *ρ* = 1.0

From the analysis of the long time dynamics of the system (29), and by taking into account different timescales of processes, we have shown that there are two fixed points where *c* = 0 or *c* = *c*\*. If the MHC-I expression is above some critical value, *γ* > *γ* _*c*_, the fixed point *c* = 0 is stable and *c* = *c*\* is unstable, whereas the reverse is true if *γ* < *γ* _*c*_. In doing so we have assumed that *τ*_*a*_ ≪ *τ*_*c*_. If we assume the reverse *τ*_*c*_≪ *τ*_*a*_, the exact trajectories will differ, however the qualitative analysis of the long-time dynamics will be the same.

This shows that there is a critical value of MHC-I expression, above which the tumour will be eradicated, and below which it will reach some stable tumour size. Furthermore, the critical value of MHC-I depends linearly through the helper/cytotoxic ratio *R*. To remove tumour cells a large population of cytotoxic T-cells is required and this depends on the expression of MHC-I. This is illustrated in figure 2 (right panel). This implies that measuring CD4^+^/CD8^+^ alone may yield incorrect understanding of tumour progression *in vivo* since it also depends on the expression of MHC-I. With a low MHC-I expression, a CD4^+^/CD8^+^ ratio that would have been considered healthy for high MHC-I expression would not lead to complete tumour eradication.

The stability of the fixed points does not depend on the T-cell activation noise, controlled by the ‘temperature’ parameter *β*^−1^ = *T*. The noise does however affect the size of the escaped tumour as shown in figure 3, obtained by solving together equations (25) and (26) numerically. An additional observation is the dependence of *γ*_*c*_ on the parameters *A* and *ρ* in (28). These parameters, respectively, represent the specificity and infiltration of T-cells in the tumour, and their relationship with *γ*_*c*_ suggests that small changes in the infiltration of T-cells can greatly affect tumour growth.

**Fig. 3.**
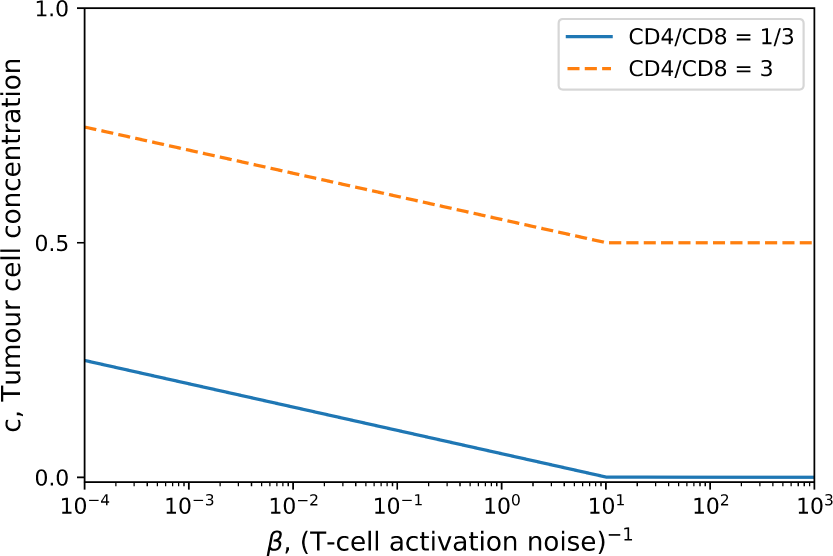
Equilibrium tumour concentration plotted as a function of inverse T-cell activation noise β. Activation noise is increasing from right to left. The tumour replication rate is *r* = 0.5, the carrying capacity concentration is *ρ*_*c*_ = 1000, and the MHC-I expression and specific tumour density are set to *γ* = *A* = *ρ* = 1. The tumour cell concentration is normalised with respect to the carrying capacity *ρ*_*c*_

### 3.2 Tumour size with immune parameters

This model provides predictions for the dependence of the size of the tumour, given by equations (25) and (26), on different immune parameters. Figure 4 shows how the size of the tumour varies with MHC-I expression (left panel) and T-cell infiltration *Aρ* (right panel) and it shows that it increases when the ratio CD4^+^/CD8^+^ is larger. For a high CD4^+^/CD8^+^ we see a discontinuity in the stable tumour size. This is due to the figure showing the tumour size dynamically reached at equilibrium. Above the critical value of infiltration, which can also be found from (28), there are enough T-cells to remove tumour cells, but below this value the tumour can grow to a stable size that depends on all other parameters of the system.

**Fig. 4.**
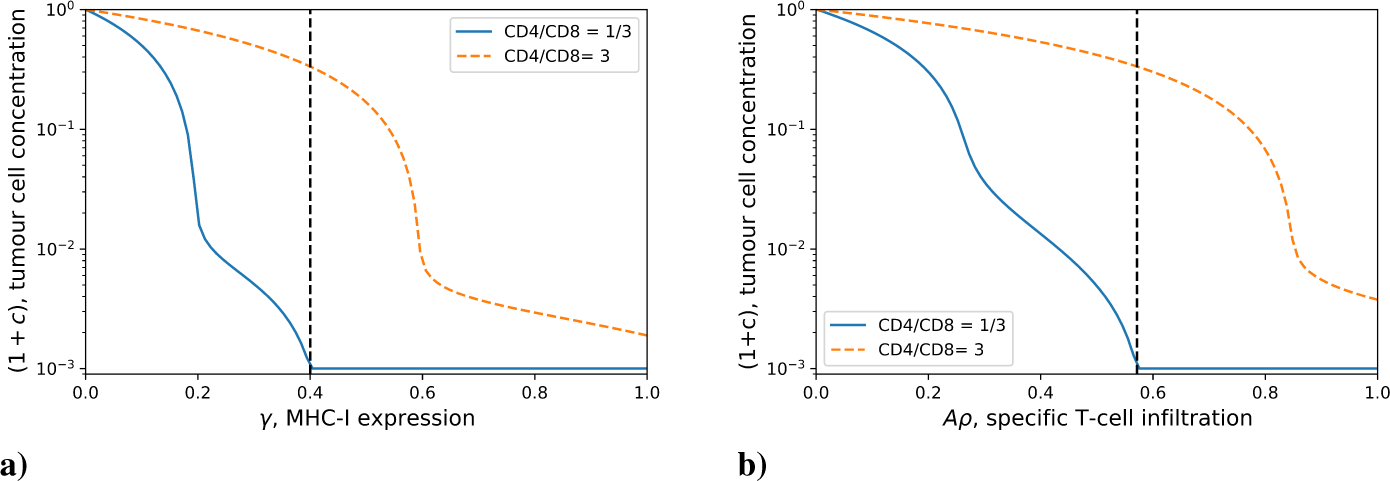
Predicted equilibrium concentration of tumour cells against plotted as a function of MHC-I expression in (**a**) and as a function of specific tumour T-cell infiltration in (**b**), displayed for two different helper/cytotoxic T-cell ratios. Here the tumour replication rate is *r* = 0.15, the activation noise is *β*^−1^ = 1, T-cell density is *ρ* = 1, the specificity in (**a**) is set to *A* = 1, the MHC-I expression in (**b**) is *γ* = 0.7, and the carrying capacity is set to *ρ*_*c*_ = 1000. The tumour cell concentration has been normalised by the carrying capacity *ρ*_*c*_. For the case of CD4^+^/CD8^+^ = 1 3 the dynamically reached equilibrium tumour cell concentration, given by equations (32) and (33), is plotted to the right and the left of the dashed (vertical) lines respectively

A common problem with ODE models of immunology is that they require temporal data for validation - which can be hard to come by for both practical and ethical reasons. Our model is based upon immune dynamics but can be used to produce predictions not based on time as in the case with the figures discussed in this section.

Concurrent measurements of MHC-I, T-cell infiltration and CD4^+^/CD8^+^ are lacking in the literature, but are examples of data which could be used to validate this model.

### 3.3 Optimal helper/cytotoxic ratio

Another feature of this model is the ability to predict an optimal helper/cytotoxic ratio *R*. Here we define optimal to mean yielding the lowest stable tumour size when equation (31) equilibrates to the fixed point *c* = *c*\*. We parameterised the system such that the fixed point *c* = *c*\* was always stable and considered the numerical solution for the stable tumour cell concentration, (25)-(27), for different values of *R* as shown in figure 5. As expected, if there are too few cytotoxic cells, the tumour will ‘grow’ to a larger size, but also if there are not enough helper cells to activate the cytotoxic population, the tumour also reaches a large size, at low noise levels, when there are no stochastic fluctuations in the cytotoxic cell activation. An interesting feature of this model is that it shows that there is a range of values of *R* for which the tumour size is relatively small. This suggests that the immune system is robust to changes in the CD4^+^/CD8^+^ ratio. This is encouraging as the function of the immune system should not be sensitive to changes in this ratio.

**Fig. 5.**
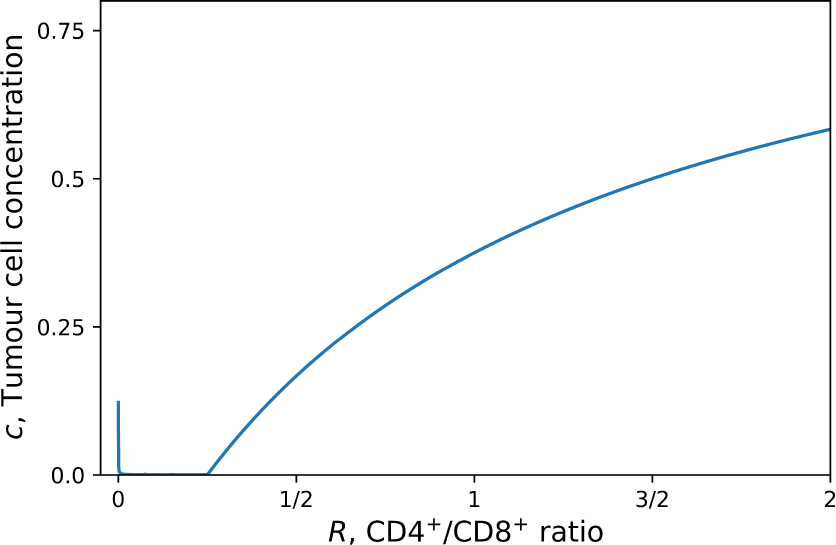
Cell concentration of a tumour which has escaped plotted as a function of the helper/cytotoxic T-cell ratio *R* for *β* = 100, *r* = 0.8, *A* = *γ* = *ρ* = 1, *ρ*_*c*_ = 1000. The tumour cell concentration has been normalised with the carrying capacity *ρ*_*c*_

### 3.4 Time variation of MHC-I expression

In our model we have treated the expression of MHC-I, *γ*, as a constant, however in principle it should evolve with time, *γ* = *γ*(*t*). As the tumour progresses, tumour cells with low expression of MHC-I will evade the immune response and will have an advantage over tumour cells that have high MHC-I expression. To understand the role of time-dependent MHC-I expression we assume a “best-case” scenario in our model such that all T-cells are active, i.e 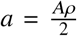 and 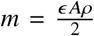, giving us the evolution of tumour cell concentration as

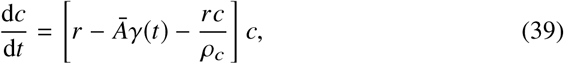

with,

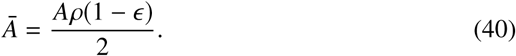

The solution of the above equation, given by

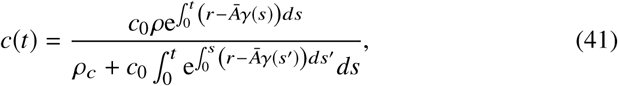

requires knowledge of *γ*(*t*) The latter has not been studied in the literature, however, we can bound the solution for a family of *γ*(*t*) functions if we assume that the maximum, *γ*_max_, and minimum, *γ*_min_, values of the function are known. We find the upper bound to be as follows

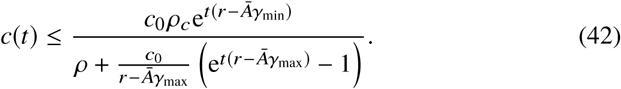

The long-time behaviour of the upper bound in the above depends on *γ*_min_ as follows

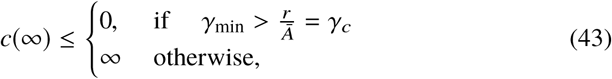

where we have recovered that for the tumour to be eradicated as *t* →∞ we require that *γ* > *γ*_*c*_.

A tighter bound on *c*(*t*) can be found with a specific form of *γ*(*t*). In particular, if we assume that the expression of MHC-I decays exponentially,

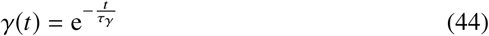

where *τ*_*γ*_ is the timescale of MHC-I decay, the solution can be bound as follows,

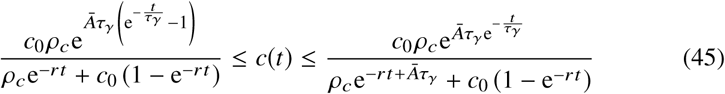

where in the above we have used 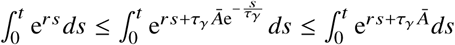. If we now consider the long-time behaviour of *c*(*t*) we find that

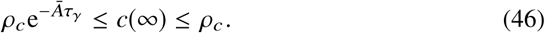

The upper bound is now finite in the long time limit, and is equal to the carrying capacity concentration, as would be expected. We see that the lower bound of *c*(*t*) is also finite. The latter is due to *γ*_min_ < *γ*_*c*_. Therefore, according to this model the exponential decay of MHC-I prohibits the eradication of tumours. However, it is important to stress that if the decay rate is sufficiently slow, the tumour cell concentration can become very small. In this regime, stochastic fluctuations, suppressed as *N* → ∞ and hence unaccounted for by our approach, become important and can remove the small amount of tumour cells. Additionally, at low MHC-I the NK cells begin to play a more dominant role, these would remove tumour cells at this point, leading to eradication.

In addition to exponential decay we have also considered a sigmoidal decay by defining

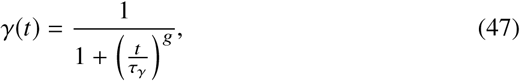

which is sigmoidal for *g* > 1 where g is the shape parameter. Furthermore, the function approaches a step function as *g* ≫ 1. Figure 6 shows that in the case where the tumour is initially growing at a rate faster than T-cell mediated death the behaviour changes significantly with the functional form of *γ*(*t*). In the case of exponential MHC-I decay, the tumour cell concentration exponentially increases towards saturation. However, with a sigmoidal *γ*(*t*), the tumour concentration increases and reaches a plateau then rapidly grows to saturation. No such difference is observed when the tumour is initially removed by the T-cells. We note that when the rate of T-cell mediated death is initially faster than the rate of replication, figure 6 exhibits the “three Es” of cancer immunoediting first discussed by Dunn et al (2004): the tumour is initially eliminated, reaches a period of equilibrium, and then escapes. This model suggests that the profile of *γ*(*t*) has a dominant effect on the duration of the elimination, equilibrium and escape phases of tumour progression. A greater quantitative understanding of the loss of MHC-I in patient tumours may reveal whether MHC-I is a dominant factor in tumour equilibrium.

**Fig. 6.**
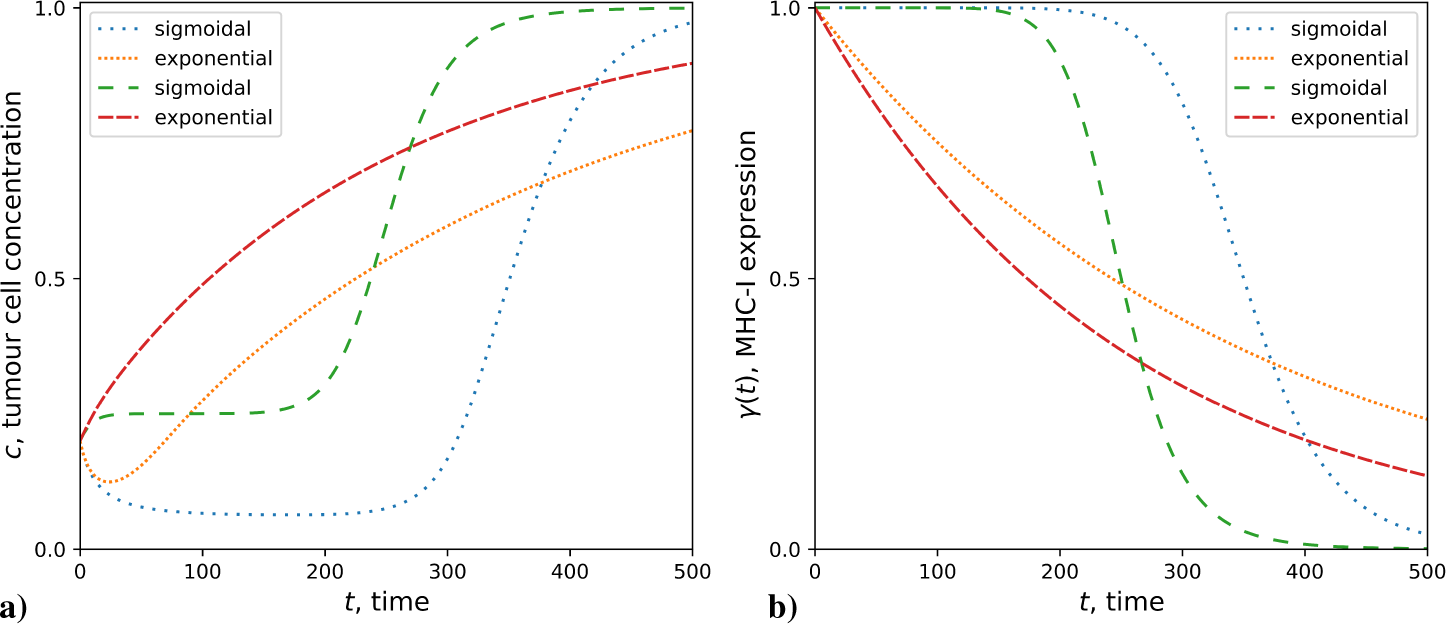
Time evolution of tumour cell concentration from numerical solution of equation (39) (**a**) subject to exponential and sigmoidal MHC-I decay profiles (**b**). We show results for two parameterisations: tumour replication rate *r* = 0.4, timescale of MHC-I expression decay *τ*_*γ*_ = 350 (dotted lines); *r* = 0.5 and *τ*_*γ*_ = 200 (dashed lines). In all cases the initial tumour cell concentration is *c*_0_ = 20, *Ā* = 0.375, and carrying capacity concentration *ρ*_*c*_ = 100. The shape parameter for the sigmoidal decay function is *g* = 10. The tumour cell concentration has been normalised with the carrying capacity *ρ*_*c*_

## 4 Discussion

The complexities of tumour immunology require a systemic approach to better understand cancer and inform treatment. Quantitative tools are being used to a greater extent, due to the vast data produced by next generation experimental technology; mathematical modelling being among them. Mathematical modelling can suffer from two extremes: either too few parameters are used, and the systemic view is lost; or there are too many parameters, which achieves greater systemic resolution, but makes statistical validation impractical, especially in absence of large data sets. In our work we have produced a model with just *five* ODEs, combining a few key parameters representing cellular processes, that capture the qualitative behaviour of the anti-tumour T-cell dependent immune response under different plausible biological conditions. To do this, we have brought over a number of techniques from non-equilibrium statistical mechanics to the modelling of biological systems.

In the past there were few works which have measured both the CD4^+^/CD8^+^ ratio and MHC-I expression together (Turcotte et al, 2014; Jordanova et al, 2008). Our hypothesis that the interplay between the two parameters may lead to contradictions in the literature is highlighted by the qualitative behaviour of our model. If the CD4^+^/CD8^+^ ratio is being used as a prognostic marker, it is clear that the expression of MHC-I must also be taken into account otherwise this may risk incorrect prognosis. A good prognosis of clinical outcome has been associated with different CD4^+^/CD8^+^ ratios across different cancers, and this model suggests that this is due to variations in MHC-I expression. If proved correct this could help to potentially unify efforts across different cancers, something that can rarely be achieved. Our model also highlights that the infiltration of specific T-cells is an important parameter, that will also affect the growth of a tumour and could potentially obfuscate the prognostic value of CD4^+^/CD8^+^ and MHC-I. Recent work has shown that the TCR repertoire of tumour infiltrating T-cells has low tumour reactivity (Scheper et al, 2019). Our model highlights the dramatic effect such a small pool of specific T-cells can have on the prognostic value of CD4^+^/CD8^+^ and MHC-I. Encouragingly, the model shows that the adaptive immune response is robust to changes in the CD4^+^/CD8^+^ ratio, as one would hope.

Our work has focused on modelling the immune response where MHC-I expression is assumed to be constant to highlight its interplay with the helper/cytotoxic ratio *R*. However, we have shown the role that MHC-I decay can play in the evolution of tumours. The eradication of tumours will depend on the minimum expression of MHC-I - if this is too low the T-cell response will fail and other cells, most likely NK, are required for eradication, otherwise the tumour will escape. In addition to this, our work highlights the role that the profile of MHC-I decay can have on the growth of tumours. To the best of our knowledge, there is no data measuring the time evolution of MHC-I expression in tumours to be able to estimate this profile. However, availability of such data could confirm whether the variation in MHC-I expression leads to periods of equilibrium in tumour growth.

One issue with mathematical models is that they often focus on the dynamics of immune cells, for which experimental data is rare. Although our model is based on dynamics, we have provided results which focus on non-temporal quantities that could be verified with standard measurements in immunology.

The work presented here is a theoretical minimal model of the adaptive immune system, and as such has limitations. For example, the model only considers the adaptive immune response and does not explicitly take into account innate immunity such as the natural killer cells and macrophages, both of which play an important role in the anti-tumour immune response. The benefit of our approach is the rigorous mathematics which can be analysed to understand qualitative behaviour, something which is often lost when considering too many parameters. Despite these limitations, the qualitative behaviour that this model produces could explain the, previously mentioned, contradictions in the literature.

A generally applicable feature of our work is the use of non-equilibrium statistical mechanics to include an additional level of microscopic detail into the modelling of cell concentrations with ODEs. Here our model has consider the activation and sub-type of T-cells, but this framework could also be used to study other biological systems where constituent cells fluctuate stochastically between different states.

Extensions to our work could look at including the heterogeneity in tumour cells. Another extension would be to adapt this model to understand how tumours begin in the first place, including the healthy tissue cells from which the tumour cells derive. Tumour immunology is rich with complexity and bringing refined tools from statistical mechanics may shed light on this complex system of processes.

## 5 Acknowledgements

All authors would like to thank Franca Fraternali and Joseph CF Ng for useful discussions regarding the direction of this work.

## 6 Declarations

### Ethics approval

N/A.

### Consent to participate

N/A.

### Consent for publication

N/A.

### Availability of data and material

N/A.

### Code availability

code is available upon request.

## A Kramers-Moyal expansion of the master equation

In this section, we derive a master equation for the probability *P*_*t*_ (***σ***) to observe a T-cell configuration ***σ*** ∈ {0, 1}^*N*^ at time *t*, from the stochastic update rule (19) and we will use it to derive equations for the time evolution of the macroscopic variables *a*(***σ***) and *m*(***σ***). Denoting by 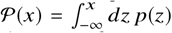 the cumulative distribution function of the noise distribution *p z*, the likelihood to observe configuration ***σ***_*i*_, at t time *t* + Δ, given the T-cell configuration ***σ***′ at the earlier time step *t*, is, for any symmetric distribution *p*(*z*) = *p*(−*z*),

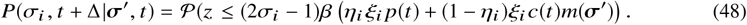

For the Glauber choice 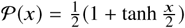, the probability that T-cell *i* changes, in a single time step, its state at time *t* is

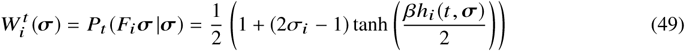

where we have defined the ‘flip’ operator *F*_*i*_ such that *F*_*i*_***σ*** = (*σ*_1_, …, 1 − *σ*_*i*_, …, *σ*_*N*_) and *h*_*i*_ (*t*, ***σ***) = *η*_*i*_ *ξ*_*i*_ *p*(*t*) + (1 − *η*_*i*_)*ξ*_*i*_ *c*(*t*)*m*(***σ***). Assuming that the update of T-cells is sequential, i.e. at each time step one T-cell *i*, drawn at random, is updated with likelihood 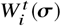, one obtains, for Δ = 1/*N* and *N* large, the following master equation

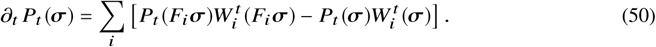

From the master equation, the time evolution of the macroscopic variables can be retrieved using the Kramers-Moyal (KM) expansion. To perform this expansion we note that we can define the time-dependent probability distribution of the macroscopic variables as

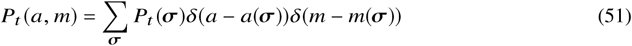

from which the master equation tells us,

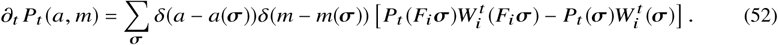

Defining **Ω**(***σ***) = (*m*(***σ***), *a*(***σ***)) and relabelling the first term in our sum with *F*_*i*_***σ*** → ***σ*** we have

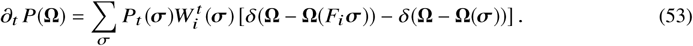

We now define the change in the macroscopic parameter Ω_*µ*_ caused by a flip in a single T-cell *i* as Δ_*iµ*_ (***σ***) = Ω_*µ*_ (*F*_*i*_***σ***) − Ω_*µ*_ (***σ***) such that 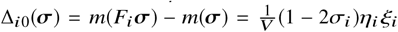 and 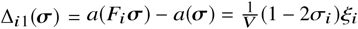. The KM expansion can then be carried out in powers of Δ_*iµ*_ (***σ***),

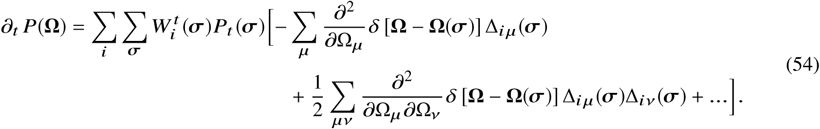

A special case where the dynamical equations close is found when 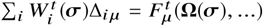, where 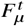 is some function that depends on the microscopic variable ***σ***, through the macroscopic variables Ω (***σ*)** only,. To this end we evaluate Σ *W*_*i*_Δ_*iµ*_ (***σ***) for the cases *µ* = 0, 1,

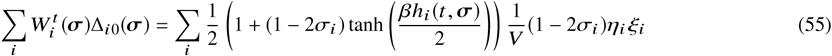

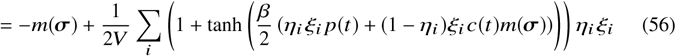

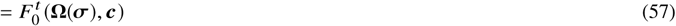

and similarly,

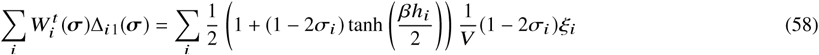

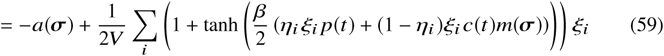

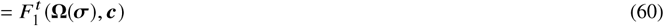

where we have denoted **c** = (*c, b, p*) and note that from equations (13)-(15), **c** only depends on ***σ*** through Ω(***σ***). Indeed, it is the case that

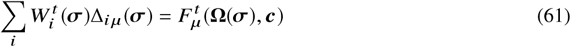

with no explicit dependence on ***σ***. By substituting equation (61) into (54) the sum over ***σ*** can be taken; this constrains Ω_*µ*_ (***σ***) = Ω_*µ*_ and yields,

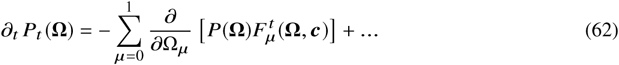

Higher order terms in the KM expansion are shown to be proportional to *V* ^−*d*^, where *d* ≥ 1, and are therefore negligible in the limit that *V* is large. Equation (62) in this limit reduces to the Liouville equation

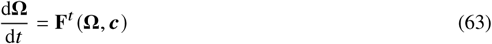

which is otherwise written,

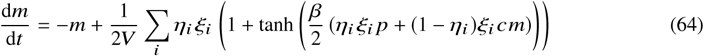

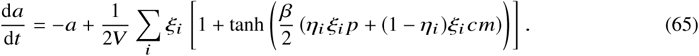

To simplify these equations further, we make use of the empirical joint distribution of *η* and *ξ*,

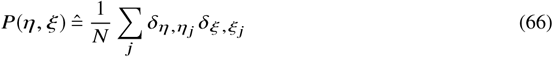

with averages over this distribution then defined as

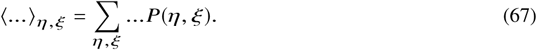

The macroscopic dynamics are then summarised by the following ODEs,

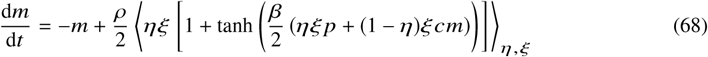

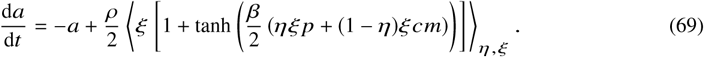

This corresponds to a mean-field description of the evolution of the stochastic variables Ω_*µ*_ (***σ***). For large but finite values of *V*, Ω_*µ*_ (***σ***) will fluctuate about its mean value Ω_*µ*_ = ⟨Ω_*µ*_ ***σ***)**⟩**_***σ***_ with fluctuations of Order 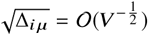

